## Supplementary figures and images for "Refined quantification of infection bottlenecks and pathogen dissemination with STAMPR"

### Figure S1

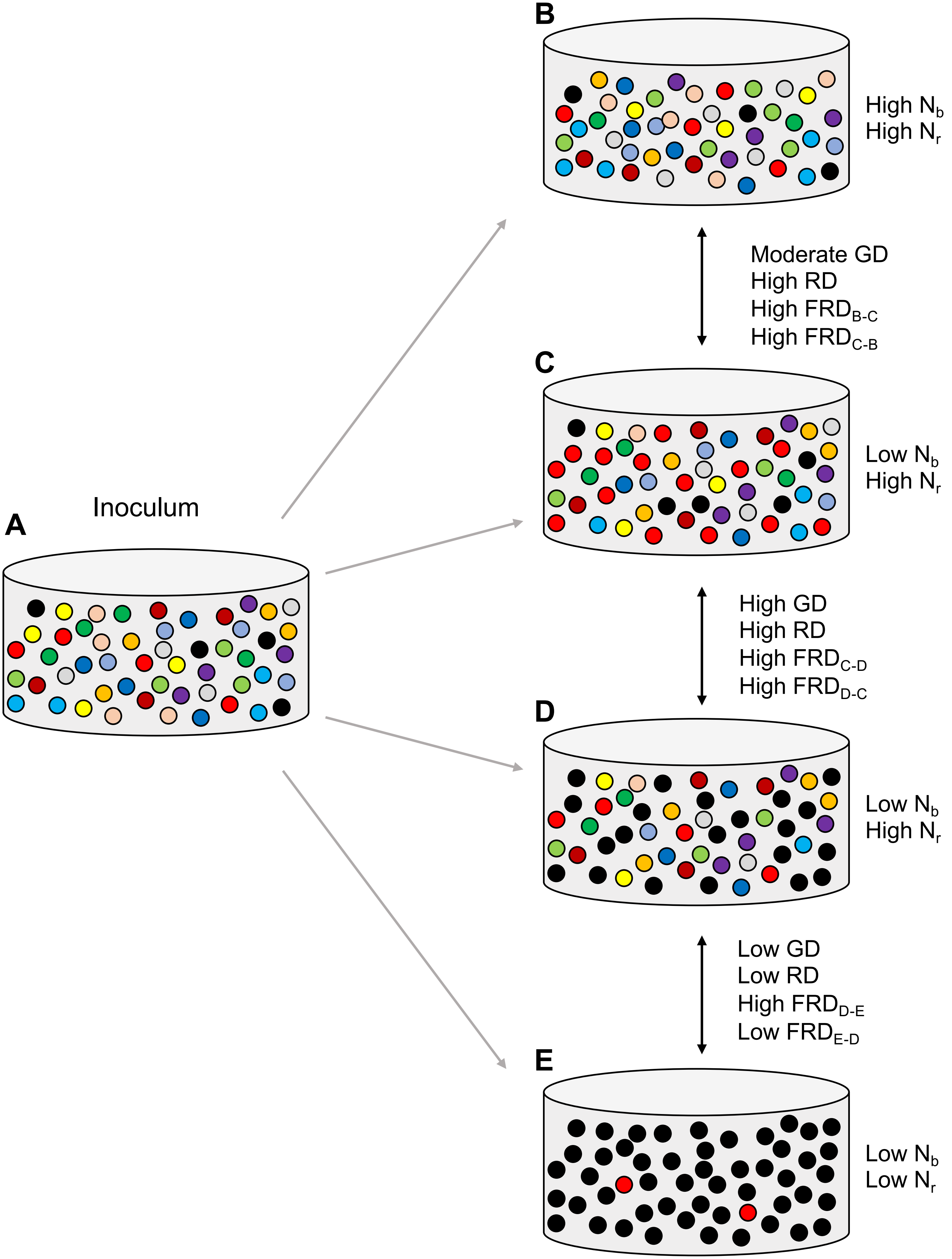

### Figure S2

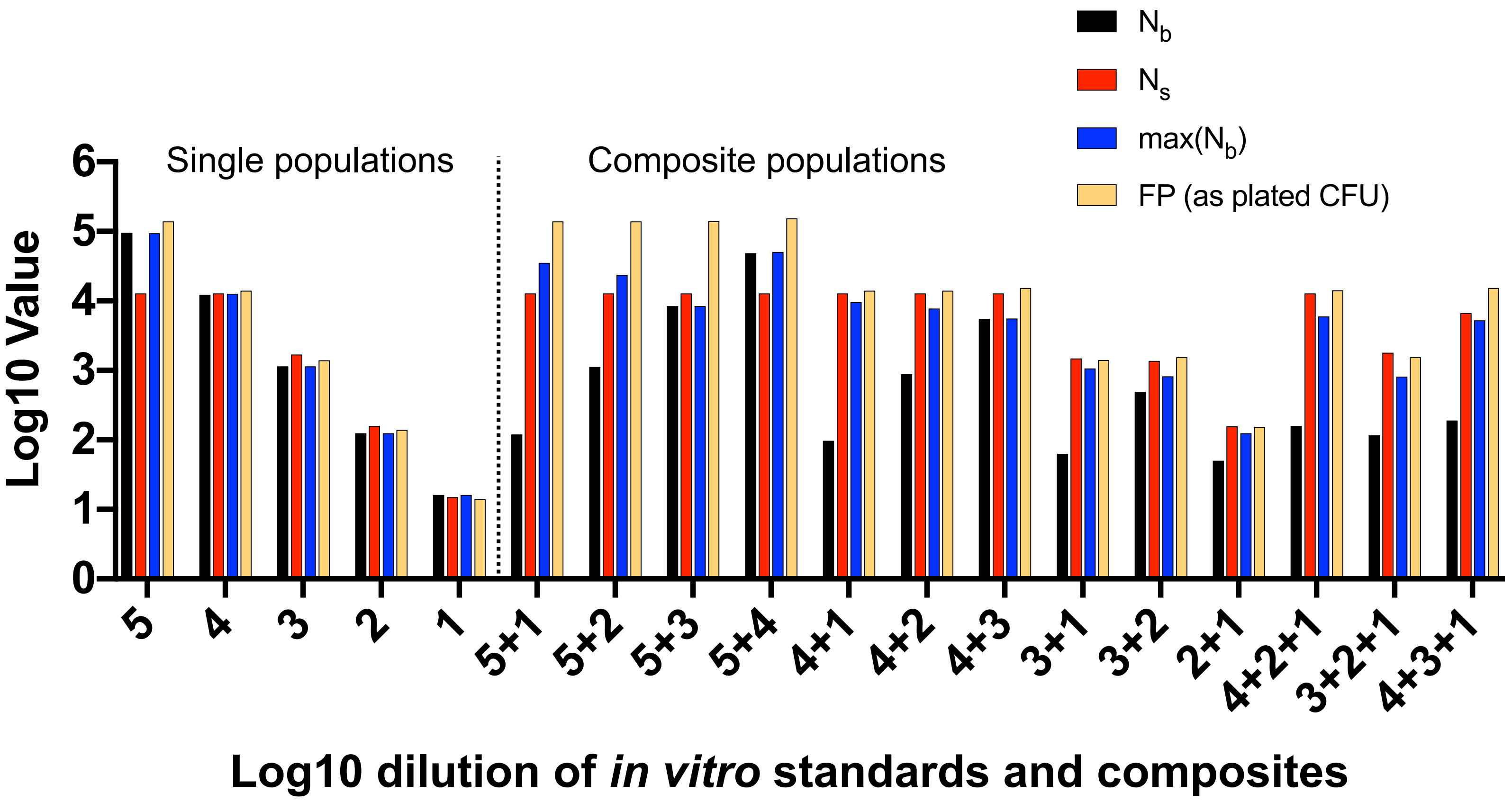

### Figure S3

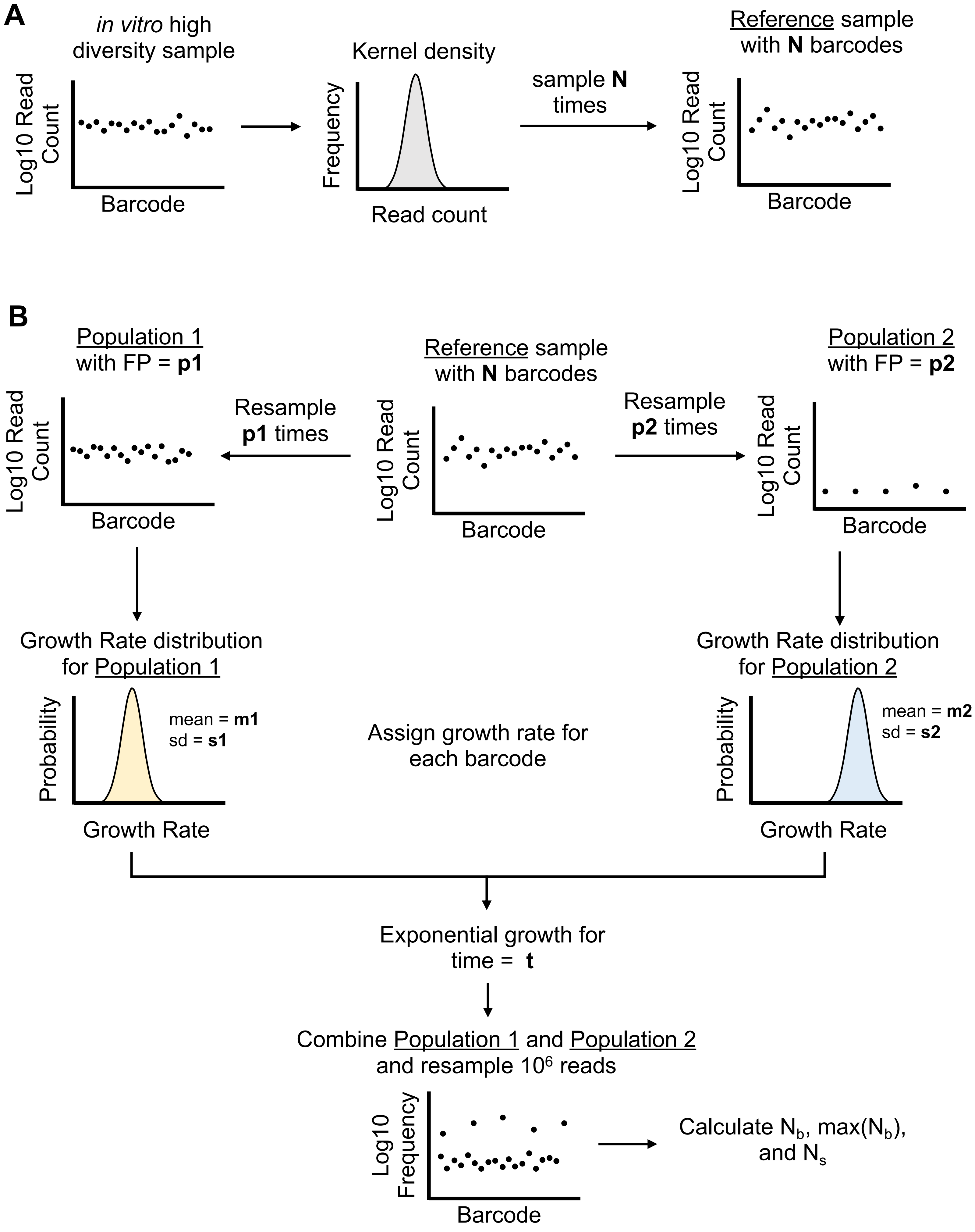

### Figure S4

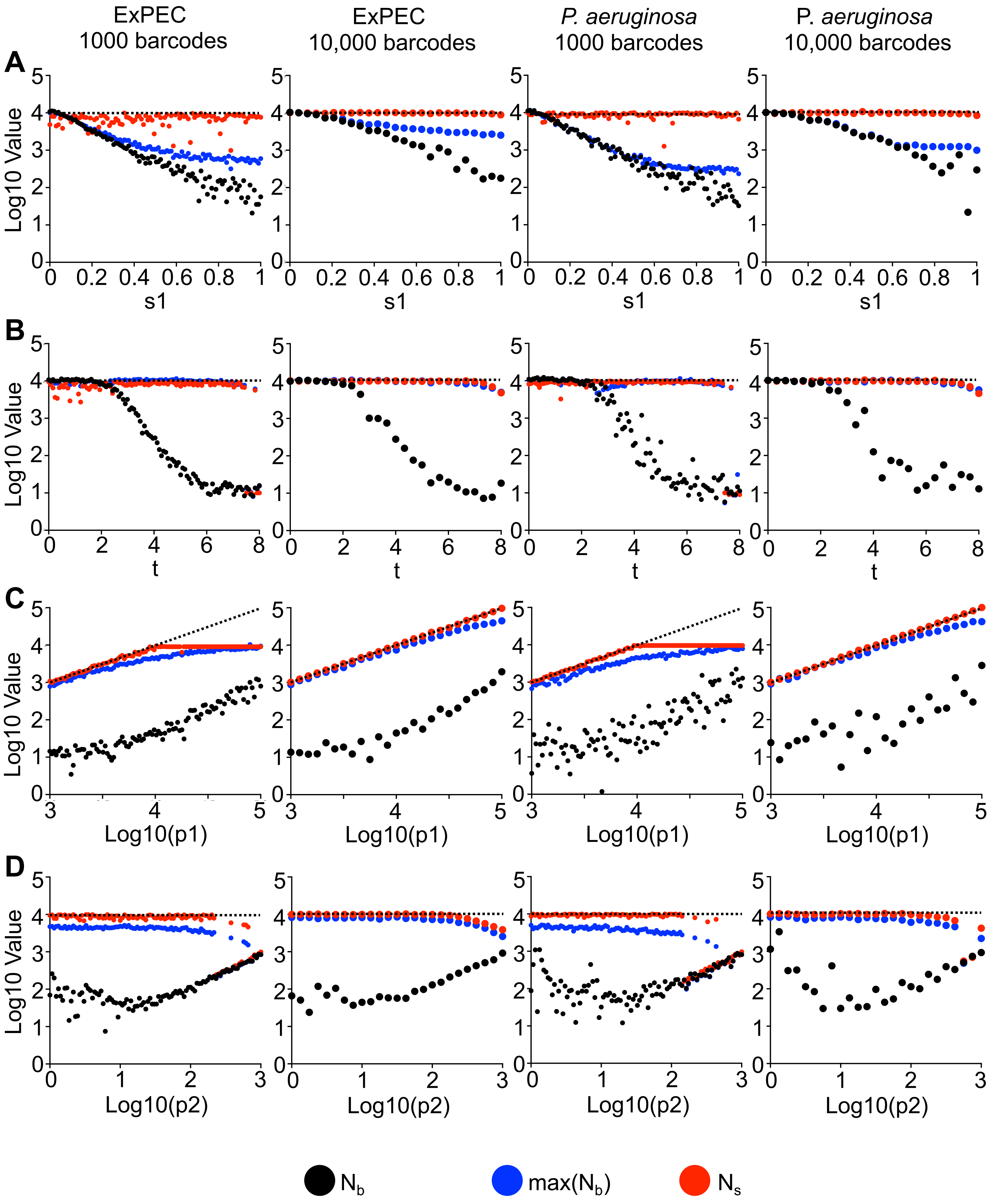

### Figure S5

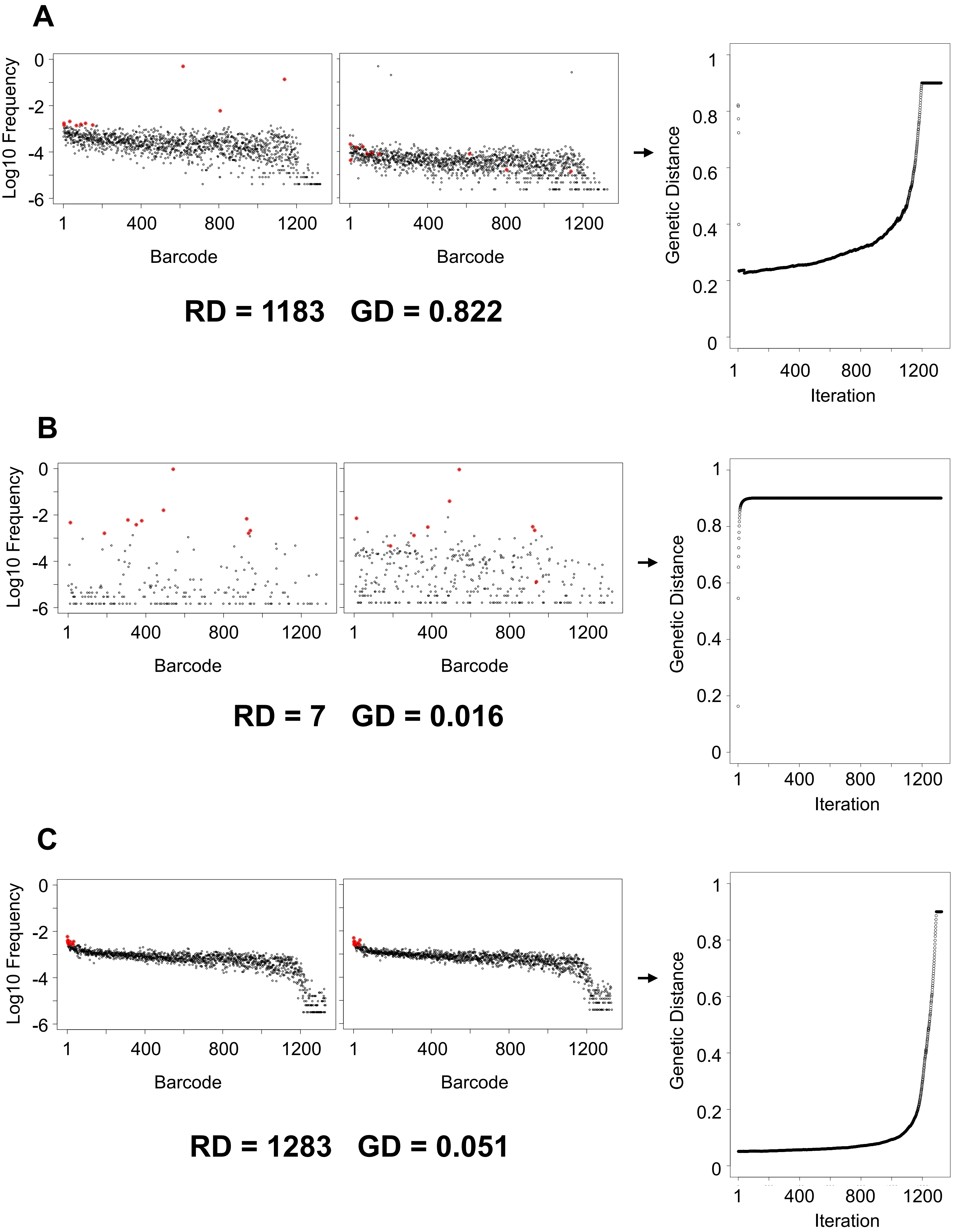

### Figure S6

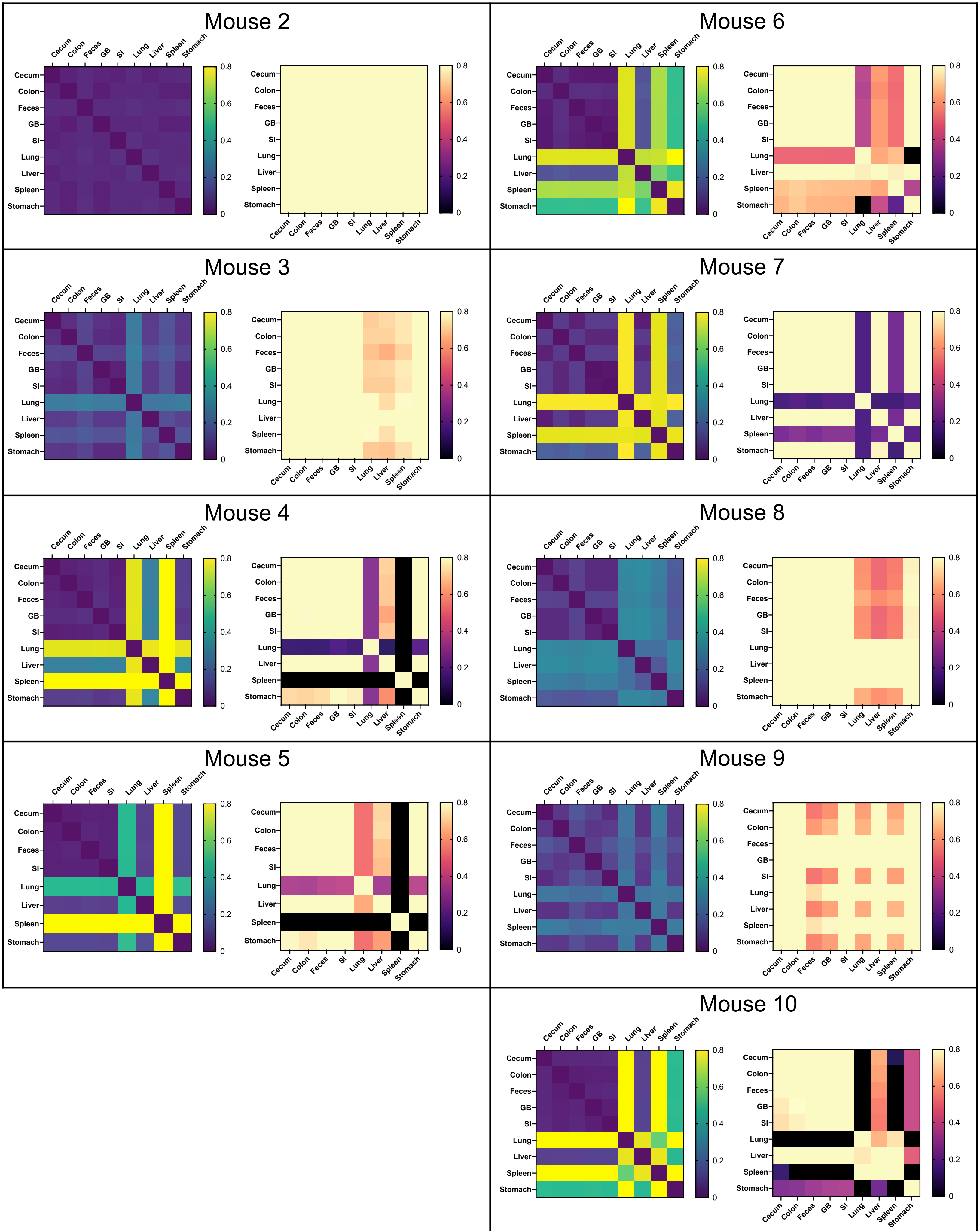

### Figure S7

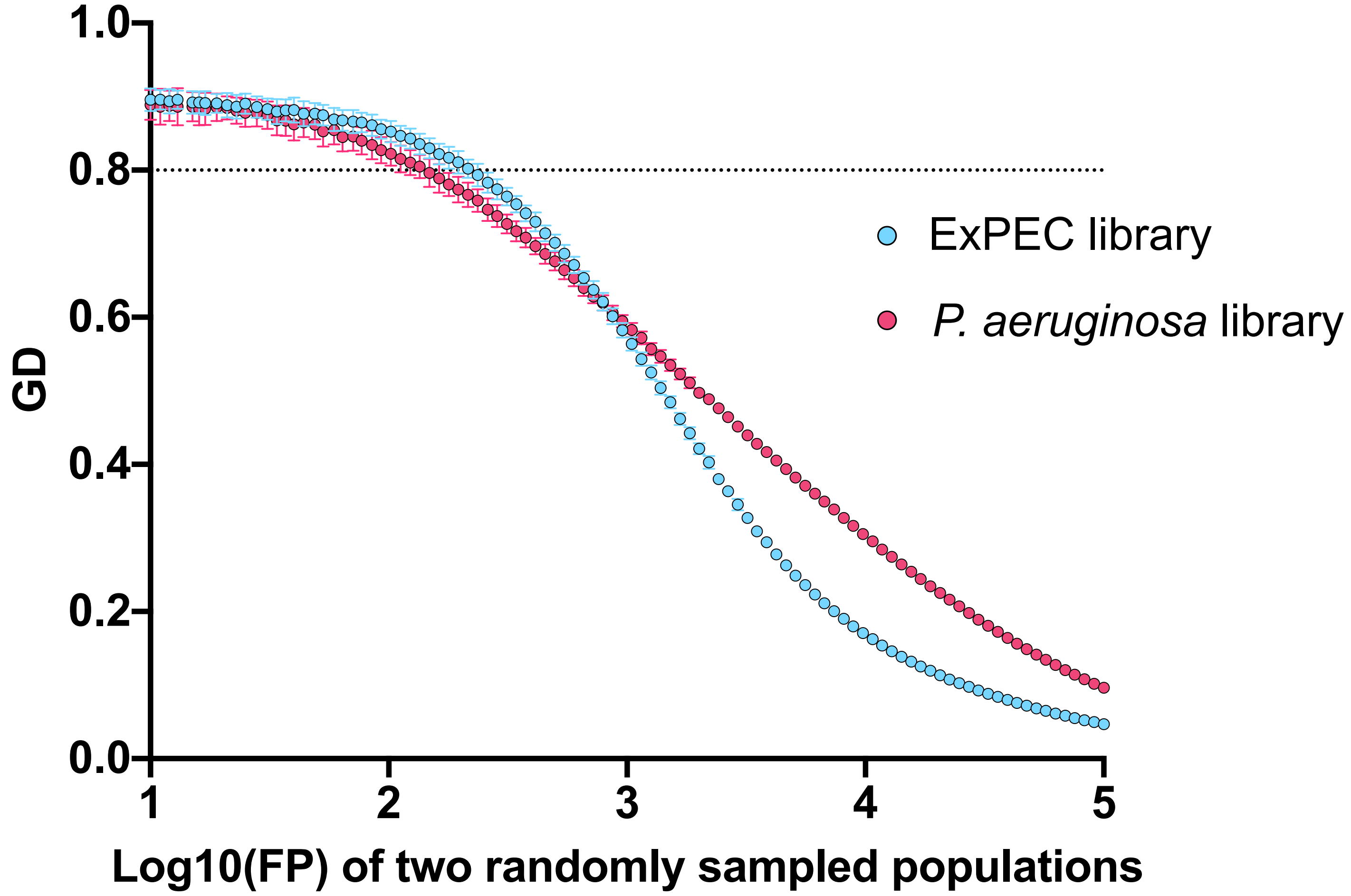
