## Supplementary material for "Refined quantification of infection bottlenecks and pathogen dissemination with STAMPR": Table S1

| **Variable** | **Description** |
| --- | --- |
| FP | The number of cells from the inoculum that give rise to the bacterial population in an organ. Cannot be directly measured. |
| N_s_ | Sampling depth of reference (inoculum) barcodes that yields the same number of unique barcodes as detected in the output (organ) sample |
| N_b_ | Estimate of FP from Krimbas and Tsakas equation (22). Decreases substantially in the presence of disproportionally abundant barcodes. Determined by the following equation, where *k* is the number of barcodes, $f_{i,s}$ is the frequency of the $i$th barcode in the sample, $f_{i,0}$ is the frequency of the $i$th barcode in the reference, $S_{s}$ is the number of reads in the sample, and $S_{0}$ is the number of reads in the reference.  $\left( \frac{1}{k}\sum_{i=1}^{k} \frac{\left( f_{i,s}-f_{i,0} \right)^{2}}{f_{i,0}\left( 1-f_{i,0} \right)} - \frac{1}{S_{0}} - \frac{1}{S_{s}} \right)^{-1}$ |
| N_r_ | Estimate of FP and output of Resiliency algorithm. Not influenced by highly abundant barcodes. Accounts for all barcodes in the sample and therefore requires noise correction |
| GD_A-B_ | Estimate of allelic similarity between two samples A and B. Low when samples are similar, high when samples are dissimilar. Equal to GD_B-A_. Calculated by the following equation, where *k* is the number of barcodes, $f_{i,A}$ is the frequency of the $i$th barcode in sample A, and $f_{i,B}$ is the frequency of the $i$th barcode in sample B.  $\frac{2\sqrt{2}}{\pi}\sqrt{1-\sum_{i=1}^{k} \sqrt{(f_{i,A}{)(f}_{i,B})}}$ |
| RD_A-B_ | Number of barcodes that contribute to GD between samples A and B. Low when two similar samples share a few highly abundant barcodes, high when two samples share many lower abundance barcodes. Equals 0 when no barcodes are shared (very dissimilar samples). Equal to RD_B-A_ |
| FRD_A-B_ | log((RD_A-B_ + 1) / log(number of nonzero barcodes in sample B + 1). FRD_A-B_ is low when RD_A-B_ is low but many barcodes are detected in sample B, consistent with the presence of many non-transferred barcodes in sample B |
