## Supplementary material for "Refined quantification of infection bottlenecks and pathogen dissemination with STAMPR": Text S1

**reference** 🡨 vector of barcode frequencies in the reference (inoculum) sample

**sample** 🡨 vector of barcode frequencies in the output (organ) sample

**cfu** 🡨 CFU of output organ sample

**inoc** 🡨 CFU of inoculum

**noisecorrect** 🡨 approximate fraction of reads known to be noise from control samples

**minimum** 🡨 fraction of reads that must be accounted for in the output

**minaccount** 🡨 1 – **minimum**

**times** 🡨 max (**cfu** + 1, number of elements in **reference** greater than 2)

\# the +1 above is useful for samples that have 1 CFU, but occasionally results in Nr values that are 1 greater than the CFU in the sample. While this is illogical, such a minor change will never have an effect on data interpretation.

**maxsteps** 🡨 10 * number of elements in **reference**

**steps** 🡨 sequence from 1 to **maxsteps** by 10

define function **getBotTable**	#Resampling-derived standard curve

input: **n**

**m** 🡨 barcode frequency distribution of multivariate hypergeometric resampling of **reference*inoc n** times

output: number of nonzero elements in **m**

**y** 🡨 **getBotTable** applied to all elements in **steps**

**u** 🡨 **NoiseCorrect** * number of reads in **sample**

**noise** 🡨 barcode frequency distribution from multivariate hypergeometric resampling of **reference*inoc u** times

**sample** 🡨 **sample** - **noise**

define function **getNb**	#Computes Nb

input: two vectors of barcode frequencies

output: N_b_ by Krimbas and Tsakas

**x** 🡨 0

**bind** 🡨 combined table with **reference** and **sample** ordered by elements in **sample**

define function **IterativeRemoval**	#Removes barcodes and calculates Nb

input: two column table

**t** 🡨 perform **getNb**

**bind** 🡨 remove bottom row from table

output: append **t** to **x**

**x** 🡨 Perform **IterativeRemoval** on **bind times** times

remove first 0 from **x**

**w** 🡨 number of elements in **x**

**q** 🡨 sequence from 1 to **w** evenly spaced (**w** / 15) times

#To initiate on more sites, increase the denominator

**p** 🡨 values of **x** at position = **q**

define function **ScanMinima**	#Identifies local minima

input: **start**

define function **FindMinima**

**location** 🡨 position where **x** = **start**

**newlocation** 🡨 random sampling of a normal distribution with mean = **location** and sd =0.1 * **w**

\# To find more local minima, decrease sd

**newvalue** 🡨 value of **x** at position = **newlocation**

if **newvalue** < **start**

**start** 🡨 **newvalue**

else **start** 🡨 **start**

output: **start**

**start** 🡨 repeat **FindMinima** 1000 times

**decision** 🡨 position of **x** where **x** = **start**

output: **decision**

**guesses** 🡨 apply **ScanMinima** to all elements in **p**

**guesses** 🡨 append **w** to **guesses**

**guesses** 🡨 set all guesses within 5 positions of **w** to be equal to **w**

**guesses** 🡨 sorted unique elements in **guesses**

**deltax** 🡨 position of **x** where log(**x**_i+1_) – log(**x**_i_) is greatest across all elements of **x**

**breaks** 🡨 append **deltax** to **guesses**

if **deltax** is within 1 of any value of **guesses**, remove the lower value

**maxNb** 🡨 maximum value of **x** until each element in breaks

**weights** 🡨fractional abundance of all elements of **sample** (ordered) within adjacent values of **breaks**

**IndicesTable** 🡨 append columns of **maxNb**, **weights**, and **breaks**

**NoiseStart** 🡨 value of **breaks** in row = max(log(**weights**_i+1_) – log(**weights**_i_)) in **IndicesTable**

if no values of **weights** are less than **minaccount**

**NoiseStart** 🡨 max(**breaks**)

if the sum of **weights** after the row in **IndicesTable** containing **NoiseStart** is greater than **MinAccount**

**threshold** 🡨 first weight in **IndicesTable** after which the cumulative sum of **weights** is greater than **minimum**

**NoiseStart** 🡨 element of **breaks** in **IndicesTable** with row containing **threshold**

**j** 🡨 **w** – **NoiseStart**

set the bottom **j** elements in **sample** to 0

**SecondBind** 🡨 append two columns of **reference** and **sample** and order by **sample**

**z** 🡨 0

define function **IterativeRemovalAfterNoise**	#Removes barcodes and calculates Nb

input: two column table

**t** 🡨 perform **getNb**

**SecondBind** 🡨 remove bottom row

output: append **t** to **z**

**SecondTimes** 🡨 number of nonzero elements in **sample**

**z** 🡨perform **IterativeRemovalAfterNoise** on **SecondBind SecondTimes** times

**Ns** 🡨 inverse interpolate x = **steps** and y = **y**. Solve x for y = **SecondTimes**

**Nr** 🡨 max (first element of **x**, max(**z**), **Ns**)
